# DeepMASS: Unknown Compound Annotation using Semantic Similarity of Mass Spectral Language and Chemical Space Localization

**DOI:** 10.1101/2024.05.30.596727

**Authors:** Hongchao Ji, Ran Du, Qinliang Dai, Meifeng Su, Yaqing Lyu, Yanchun Peng, Jianbin Yan

**Affiliations:** Shenzhen Branch, Guangdong Laboratory for Lingnan Modern Agriculture, Key Laboratory of Synthetic Biology, Genome Analysis Laboratory, Ministry of Agriculture and Rural Affairs, Agricultural Genomics Institute at Shenzhen, Chinese Academy of Agricultural Sciences, Shenzhen, 518120, China PR

## Abstract

Untargeted analysis using liquid chromatography□mass spectrometry (LC-MS) allows quantification of known and unknown compounds within biological systems. However, in practical analysis of complex biological system, the majority of compounds often remain unidentified. Here, we developed a novel deep learning-based compound annotation approach via semantic similarity analysis of mass spectral language. This approach enables the prediction of structurally related compounds for unknowns. By considering the chemical space, these structurally related compounds provide valuable information about the potential location of the unknown compounds and assist in ranking candidates obtained from molecular structure databases. Validated with two independent benchmark datasets obtained by chemical standards, our method has consistently demonstrated superior performance compared to existing compound annotation methods. A case study of the tomato ripening process indicates that DeepMASS has significant potential for metabolic biomarker identification in real biological systems. Overall, the presented method shows considerable promise in annotating metabolites, particularly in revealing the “dark matter” in untargeted analysis.

---

Mass spectrometry (MS) plays a crucial role in the analysis of the diverse small-molecule content of complex systems, particularly in life omics. It significantly contributes to chemobiological research, encompassing areas such as the metabolome[1], biosynthesis[2], foodomics[3], exposome[4], and. However, despite the widespread application of this technology, the structural annotation of unknown compounds remains a pressing challenge. The most commonly used strategy for structural identification is to match the detected compound MS/MS spectra with standard spectral libraries such as GNPS, MassBank and NIST. However, this strategy is limited in two ways: first, different instrument types, conditions, and parameter settings can result in the same compound having different MS/MS spectra, reducing the universality of standard spectral libraries and consequently decreasing the identification of known molecules in practical research[5]. Second, the coverage of standard spectral libraries is much smaller than the potential number of molecules, and the absence of standard spectra makes it nearly impossible to identify novel compound through library matching. These unidentified compounds become neglected “dark matter” in research.

Improvements in library-searching algorithms, such as MS-Entropy and MS2Query enhanced the robustness of spectral matching, partially solves the first challenge and improve the efficacy and efficiency of molecular annotation[6–8]. However, the “dark matter” problem remains unaddressed. For example, over 90% of chemical signatures remaining uncharacterized in untargeted plant metabolomics studies[9]. Although this bottleneck can be partially alleviated by expanding the number of standard spectra, molecules that are easy to obtain as standards are usually already included, while those not included are often difficult to synthesize and isolate. Consequently, adding new standard spectra entails higher costs and greater effort. Molecular networking, on the other hand, can reveal the chemical neighbors of unknown molecules and estimate their class, but it cannot predict their specific structures automatically[10–12].

Therefore, employing computational methods predicting chemical structures from the spectra excluding from spectral database is the most promising approach for improving coverage[13,14]. Currently, the published methods can be mainly divided into in silico annotation algorithms[15–20] and known-to-unknown extension through pathways[21–23]. In-silico annotation algorithms offer flexible application to any source and any number of unknown spectra. However, compounds that can be annotated through known-to-unknown extension must originate from metabolites that have already been annotated through spectral matching. In-silico annotation algorithms typically involve two steps. First, they find candidates from structural database[24] based on the chemical formula. The chemical formula can be inferred from the precursor masses, ionization types and isotopic pattern, or through specialized toolkits such as BUDDY[25] or MIST-CF[26]. Second, they assess and prioritize potential candidates based on specific rules. For example, SIRIUS scores candidates by comparing the predicted fingerprints of an unknown compound with the calculated fingerprints of potential candidates[16]. MS-Finder scores candidates by considering the frequency of hydrogen rearrangement inferred from the fragment ions[17]. CFM-ID evaluates candidate compounds based on the accessibility of the inferred substructural motif from the fragment ion[18].

Previous study revealed that structurally related compounds would exhibit similarities in their MS/MS spectra and deep neural networks are able to learn this correlation[27–30]. Particularly, the DeepMASS structural-similarity scoring model is proposed and verified this finding can be applied for compound identification with MS/MS, as a set of predicted structurally related compounds can suggest the possible structure of the unknown compound. However, the original work only serves as a conceptual prior study. Because it relied on constructing compound pairs and using fully connected neural networks to predict structural similarity based on their binned spectra, the model could only be trained hundreds of spectra in the training set due to combinatorial explosion. This limitation prevented it from covering a wide range of chemical spaces, seriously affecting its applicability.

We here introduce DeepMASS v2, a subversive iteration, which alternates the structural-similarity scoring strategy to an unsupervised embedding method. This enhancement enables over three orders of magnitude of the training set, making its range of applications broad enough to cover various areas of chemobiological research. Furthermore, the whole algorithm was rewritten, optimized and integrated as a GUI software, providing a user-friendly interface that allows researchers to easily utilize its capabilities for compound identification and structural inference in their studies. We have demonstrated its annotation performance by comparing it with competitive tools using benchmark datasets. Additionally, we applied DeepMASS in a specific context, showcasing its utility in assisting plant metabolomics studies.

## RESULTS

### Workflow and principles of DeepMASS

DeepMASS is situated within the MS/MS data process following the data preprocessing of raw files. Its objective is to accurately annotate compounds, regardless of their inclusion in the reference library. At this stage, molecular features along with their corresponding mass spectra are exported by data preprocessing software such as XCMS[31], MZMine[32], OpenMS[33], KIPC2[34] or MS-DIAL[35]. DeepMASS initiates by accepting the spectrum of features and proceeds to retrieve structural candidates of the unknown compounds from a comprehensive structural database. If the chemical formula has already been assigned, the retrieval process is based on the chemical formula. Otherwise, it alternatively retrieves candidates based on the monoisotopic mass derived from precursor masses and ionization types. All candidates with a monoisotopic mass difference within a specific threshold (10 ppm as default) are considered. The core task then becomes ranking the candidates (**Fig. 1**).

**Fig. 1.**
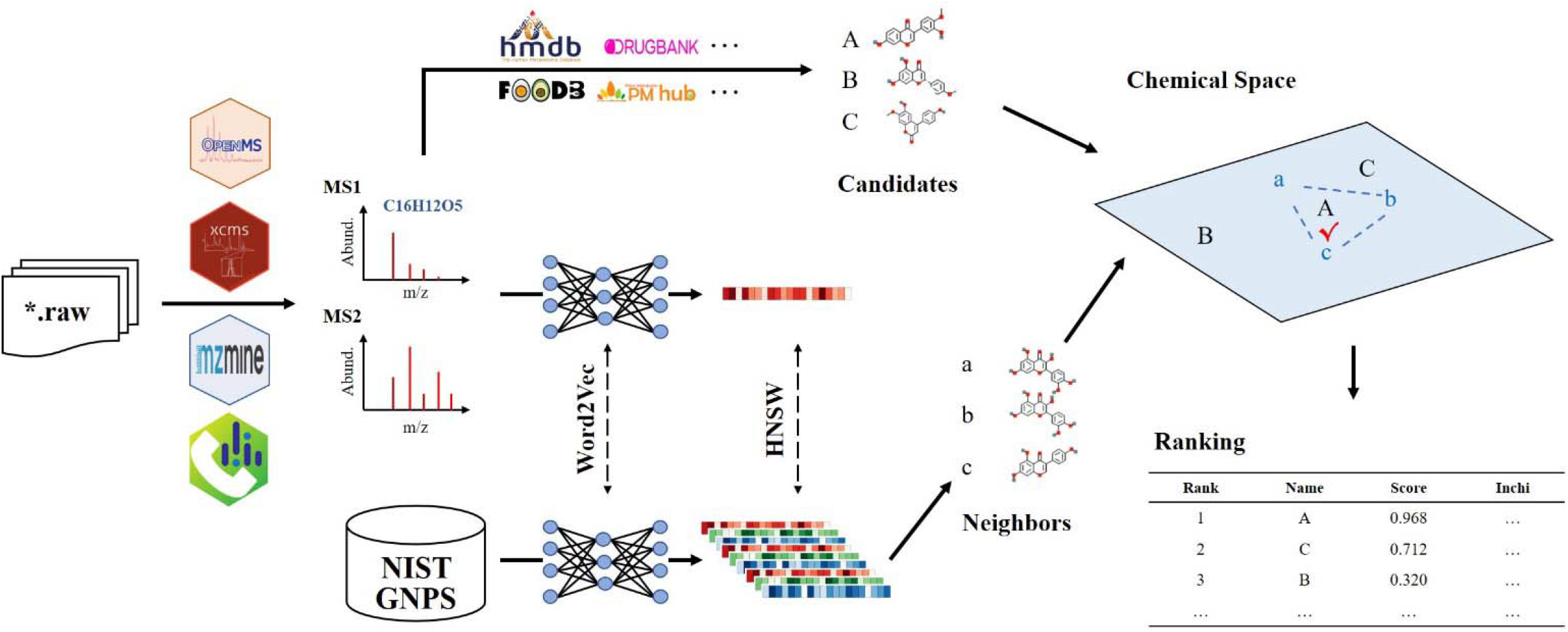
Workflow of DeepMASS. Raw mass spectrometry data are preprocessed, and unannotated compound signals are parsed into MS1 and MS2 spectral signals. MS1 is employed to predict the molecular formula, which is then used to retrieve candidate structures from chemical libraries. MS2 is embedded and used to search for structurally similar neighbors using a semantic embedding model trained on existing spectral databases. Both candidate and reference structures are projected into chemical space based on chemical fingerprints. The correct structure of the unknown spectrum is determined based on the relative positions in the chemical space of the reference structures.

The principle of DeepMASS is to identify structurally related compounds from the reference library and determine the most likely structures of the unknown compound from the candidates based on these relationships. Firstly, it trains a Word2Vec-based semantic similarity model embedding the unknown spectrum into a numeric vector. Word2Vec is widely used for comparing semantic similarity in natural language processing, which can facilitate capturing the nuanced relationships between words. Similarly, spectral features are related to each other by neutral loss, so they are also supposed to have comparable relationships to the word embedding. This strategy was applied by the Spec2Vec package[28,29], which is designed for embedding spectra for library matching and molecular networking. It has been demonstrated that the correlation between the embedded vectors exhibits a stronger relationship with structural similarity compared to cosine-based scores. Therefore, DeepMASS adopts the trained model transform both the queried spectrum and the spectra in reference library into vectors for a quick searching in the next step.

Then, a rapid hierarchical navigable small-world graph (HNSW)[36]-based search is conducted between the vectors transformed from the queried spectrum and the spectra in reference library. This aims to find the nearest neighbors of the unknown compound from the reference library. All the neighbor compounds encompass the potential chemical space of the unknown compound. Their spatial positions in the chemical space are determined using Morgan molecular fingerprints, which provide a detailed representation of molecular structure. The relative distance between compounds in the chemical space reflects their structural similarity.

Finally, the retrieved candidates are compared with the predicted neighbor compound in the chemical space. The relative spatial distances between a candidate and the predicted neighbors are measured based on the Dice similarity of the fingerprint vectors. It is expected that the correct candidate for the unknown compound will be surrounded by several predicted neighbor compounds. Based on this assumption, a similarity scoring function is employed to rank the candidates, aiding in the annotation of the unknowns.

### Construction of comprehensive database

To provide a comprehensive database for retrieving structural candidates across different research areas, DeepMASS collects, organizes, and integrates chemical structures from various databases (**Table S1**). This extensive database comprise 18 different sources, including ChEBI[37], BloodExp[38], DrugBank[39], ECMDB[40], FooDB[41], HMDB[42], KEGG[43], NANPDB[44], NPAtlas[45], PMHub[46], PMN[47], SMPDB[48], STOFF[49], T3DB[50], TCMSP[51], YMDB[52], GNPS[53] and NIST[54]. These sources cover a wide range of possible chemical structures derived from humans, plants, microbes, foods, herbs, drugs, toxicants and the environment, ensuring broad applicability across various fields of chemobiological research. As a result, 559,589 unique compounds are included. Their chemical classification are predicted with NPClassifier[55].

### Benchmark test with merged CASMI dataset

We assembled a benchmarking dataset comprising 499 spectra associated with 478 unique compounds obtained from the Critical Assessment of Small Molecule Identification (CASMI) for the years 2014, 2016, and 2022 (**Table S2**). In order to prevent data leakage, we have excluded all spectra from the training data that exhibit a cosine similarity exceeding 0.95 when compared to any spectrum in this benchmarking dataset. Statistically, 217 out of the 499 spectra correspond to compounds included in the reference dataset, setting the upper limit of spectral matching at 43.4%. Meanwhile, 455 out of 499 spectra, along with their corresponding compounds, are present in the bio-database, establishing the theoretical upper limit of in silico annotation performance at 91.2%.

Firstly, we evaluated the annotation accuracy of DeepMASS by directly comparing with the state-of-art in silico compound annotation software tools SIRIUS[16] and MS-Finder[19]. Additionally, we compared it with spectral matching using Spec2Vec to demonstrate how DeepMASS improves annotation accuracy through the integration of chemical space location, not just through the semantic embedding model. The compound databases utilized by SIRIUS and MS-Finder were the same as those used by DeepMASS, through custom uploads via their parameter setting functions. Meanwhile, we also evaluated the public version of DeepMASS, which excludes NIST spectra from the reference dataset and is available to all users. As shown in **Fig. 2a**, among all the test spectra, DeepMASS attained a Top 1 accuracy of 57.7%, signifying that it correctly placed 288 true annotations in the top position, in contrast to 44.7% for SIRIUS, 36.1% for MS-Finder, and 38.9% for Spec2Vec-based library matching. The public version of DeepMASS also achieved a Top 1 accuracy of 52.1%, still superior to both SIRIUS and MS-Finder. Upon expanding the evaluation to the Top 10 annotations, DeepMASS, DeepMASS (public), SIRIUS, and MS-Finder exhibited improved performance, with annotation rates of 79.6%, 77.8%, 65.8%, and 68.1%, respectively. However, the improvement in the performance of library matching methods were not significant because a large portion of the compounds corresponding to the test spectra were not included in its reference database.

**Fig. 2.**
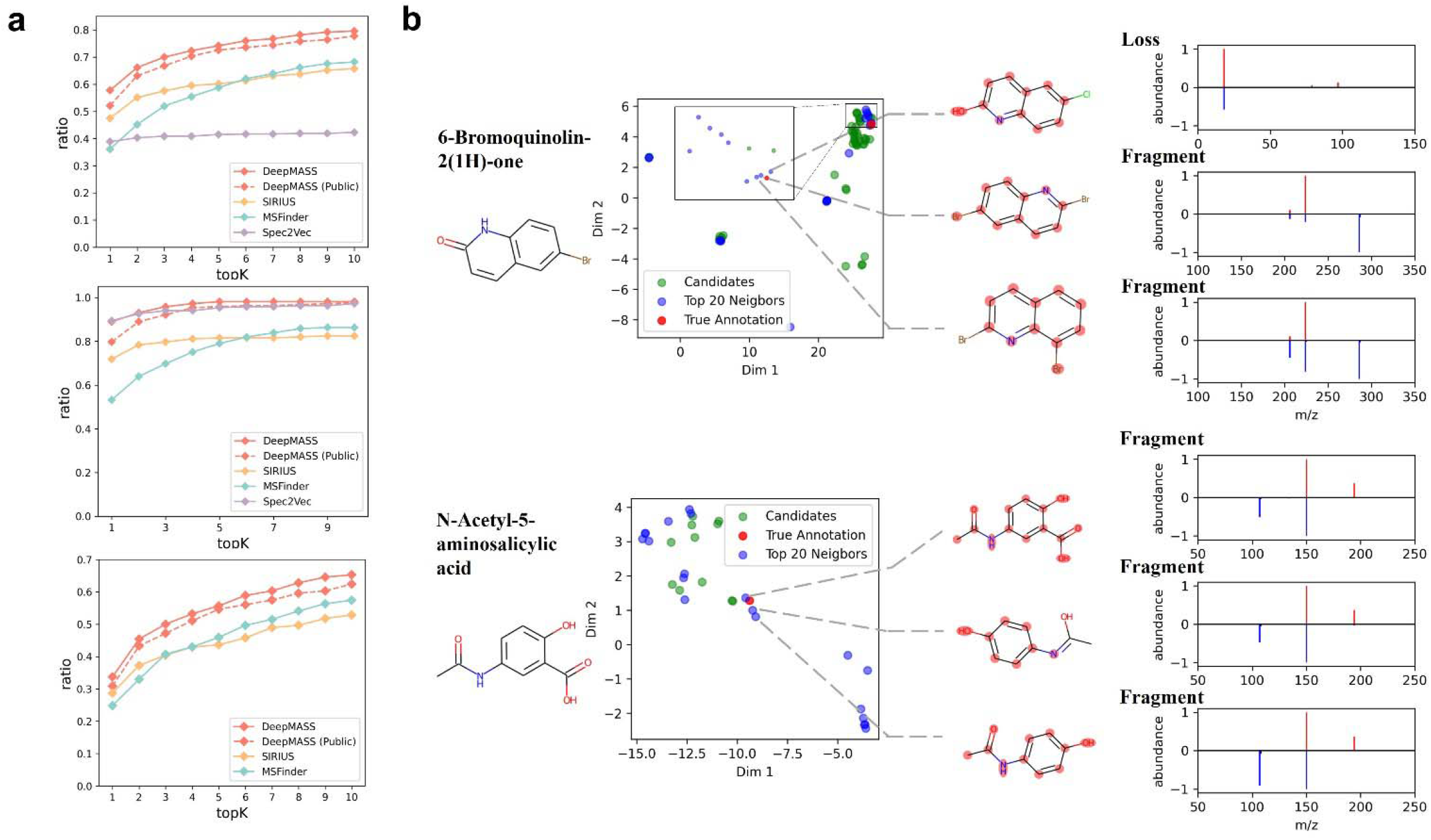
Performance of DeepMASS on CASMI dataset. **a**. Ratio of correctly identified structures found in the top *k* outputs of the different methods. The three-line plots (from top to bottom) represent the performance of all spectra, spectra with corresponding compounds present in the reference database, and spectra with corresponding compounds absent from the reference database. **b**. Annotation example for 6-bromoquinolin-2(1H)-one (absent from reference dataset) and N-acetyl-5-aminosalicylic acid (included in the reference dataset). The scatter plots are visualized in chemical space by UMAP dimensionality reduction of Morgan fingerprints. In the scatter plots, the top 10 reference structures predicted by DeepMASS’s semantic similarity model are represented as blue points, while the retrieved candidate structures are represented as green points. Notably, the correct annotations, tagged with red points, are encompassed by the predicted neighbor structures. The three closest neighbors are displayed, and shared substructures with the correct structures are highlighted. Obviously, their chemical structures are highly relevant. Meanwhile, their corresponding spectra also exhibit a high degree of correlation (red: query spectrum; blue: neighbor spectrum; fragment: the original spectrum; loss: neutral losses calculated from the precursors and fragments).

Secondly, we separately assessed the annotation performance of the spectra when matched compounds were present inside and outside the reference data. Out of the 217 spectra that included compounds from the reference dataset, Spec2Vec-based library matching demonstrated superior performance over the in-silico annotation software tools, achieving a Top 1 accuracy of 89.4%. DeepMASS achieved very similar performance, with a Top 1 accuracy of 89.0%. SIRIUS and MS-Finder attained Top 1 accuracy rates of 75.1% and 70.3%, respectively, which are also higher compared with the compounds absent from the reference dataset. This can be attributed to the fact that these compounds are more prevalent and easily identifiable by their inherent characteristics. For the compounds not included in the reference dataset, library matching was unable to provide correct annotations for any of them. In contrast, DeepMASS and DeepMASS (public) achieved Top 1 accuracy of 33.7 % and 30.9%. SIRIUS and MS-Finder yielded Top 1 accuracies of 28.7% and 24.8%, respectively. For Top 10 accuracy, DeepMASS, DeepMASS (public), SIRIUS, and MS-Finder exhibited improved performance, with annotation rates of 65.2%, 62.4%, 52.8%, and 57.4%, respectively.

Finally, we demonstrated the interpretability of the approach through two example of annotation results. They can illustrate how DeepMASS arrives at its annotation decisions (**Fig. 2b)**. We employ UMAP dimensionality reduction[56] on the Morgan fingerprints of both candidate and neighbor compounds to visualize their relationships within the chemical space. In the scatter plots, at least three of the Top 10 neighbor structures are situated near the correct annotation. These neighbor structures play a significant role in locating the chemical space of the spectrum to be annotated, thereby assisting DeepMASS in making the decision. These neighbor structures are subsequently compared to the correct structure, and their shared substructures are highlighted with a red background color.

In the case of the 6-bromoquinolin-2(1H)-one spectrum in the positive mode, the first neighbor spectrum corresponds to 6-chloroquinolin-2-ol, which exhibits losses similar to those of 6-bromoquinolin-2(1H)-one. The other two neighbor spectra correspond to 2,6-dibromoquinoline and 1,4-dibromoisoquinoline. These compounds share a highly similar carbon skeleton with 6-bromoquinolin-2(1H)-one, leading to the presence of similar spectral fragments. In the case of the N-acetyl-5-aminosalicylic acid spectrum in negative mode, the top two references originate from spectra with identical structures, resulting in duplicate scatter points with the true annotation. Nevertheless, additional references serve to further validate the position in the chemical space, specifically acetaminophen and paracetamol. Both of these references share a significant portion of the same substructure as N-acetyl-5-aminosalicylic acid. These additional references show why DeepMASS performs even better than spectrum matching when annotating spectra present in the database. Obviously, the decision-making process of DeepMASS is comprehensible and aligns consistently with theoretical expectations. The utilization of neighbor compounds predicted by the semantic similarity of mass spectral language is effective in positioning the chemical space of unannotated compounds.

### Benchmark test with natural product dataset

In order to further evaluate the performance of DeepMASS on complex compounds, we collected additional natural product benchmarking datasets. Firstly, we collected chemical standards comprising 250 known antitumor natural products and prepared five standard mixtures, each containing 50 compounds (**Table S3**). All mixtures were analyzed using a Q-Exactive Orbitrap mass spectrometer in positive ion mode. MS/MS spectra were extracted using the selected window of the calculated M+H precursor m/z. Out of the standards, 154 were successfully extracted. We evaluated the annotation accuracy of DeepMASS, SIRIUS, and MS-Finder using the same method as described above.

DeepMASS achieved the highest overall accuracy, with Top1 and Top10 scores of 47.4% and 77.3%, respectively. The public version of DeepMASS demonstrated similar performance to SIRIUS. There Top1 scores are 36.3% and 39.6%, and Top10 scores are 74.6% and 72.7%. On the other hand, Top1 and Top10 scores of MS-Finder are 7.80% and 41.1%, respectively. The declining accuracies of MS-Finder may be due to the difficulty in predicting rearrangement patterns in natural products with complex structures, as MS-Finder mainly depends on the prediction of rearrangement rules. Of the 154 spectra, 117 could be annotated using spectral matching with a cosine similarity above 0.3. For these spectra, the Top1 accuracies of DeepMASS, DeepMASS public, SIRIUS, and MS-Finder were 56.4%, 43.7%, 47.9%, and 8.55%, respectively, while greedy cosine searching achieved a Top1 accuracy of 68.4%. The Top10 accuracies for DeepMASS, DeepMASS public, SIRIUS, and MS-Finder were 86.3%, 83.8%, 80.3%, and 52.1%, respectively. For the remaining 37 compounds that could not be annotated with spectral matching, the Top1 accuracies of DeepMASS, DeepMASS public, SIRIUS, and MS-Finder were 18.9%, 13.5%, 13.5%, and 8.00%, respectively, with corresponding Top10 accuracies of 48.6%, 45.9%, 48.6%, and 40.0%. Overall, DeepMASS demonstrates similar or better accuracy compared to competitive tools in both cases (**Fig. 3a)**.

**Fig. 3.**
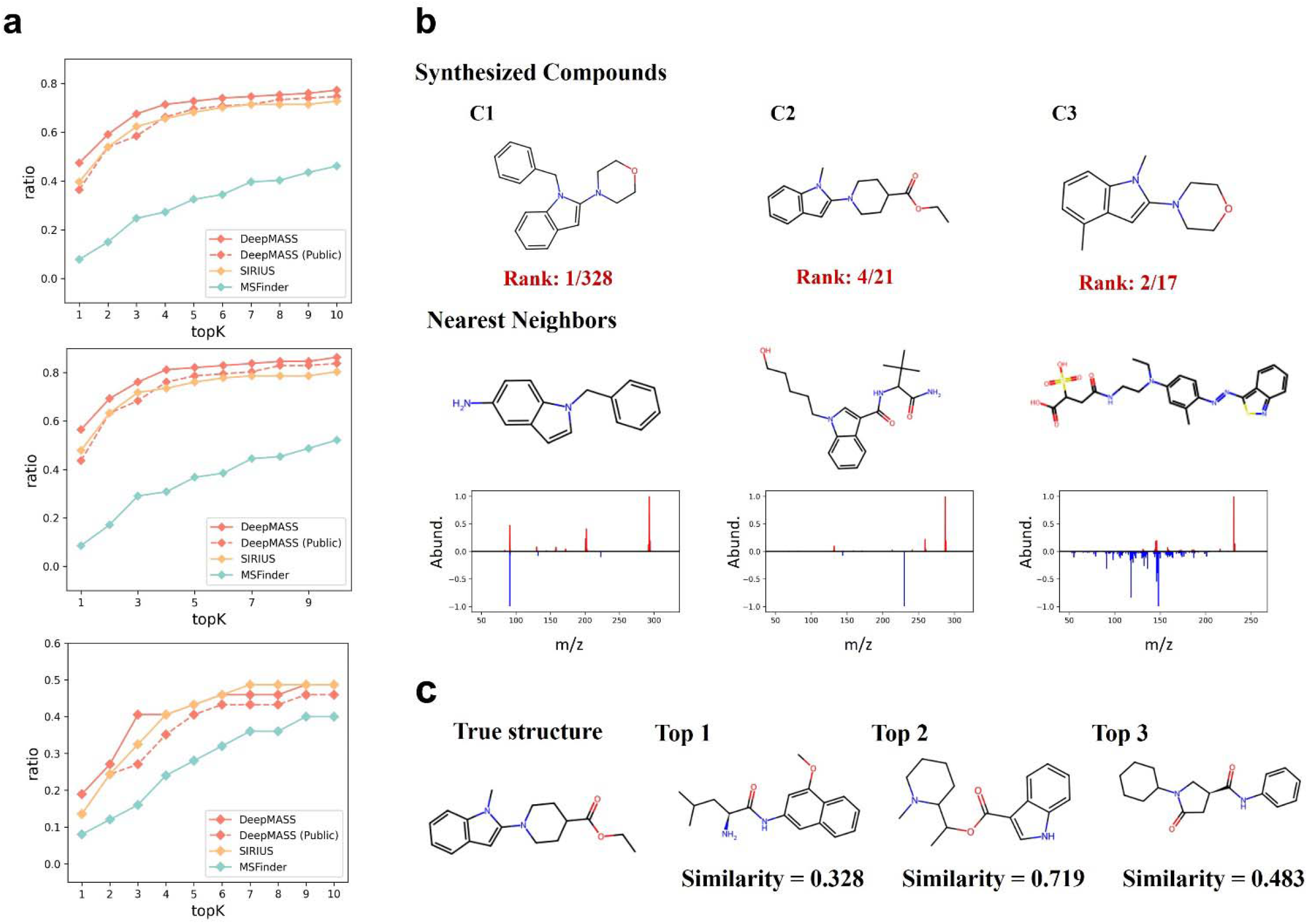
Performance of DeepMASS on natural product dataset. **a**. Ratio of correctly identified structures found in the top *k* outputs of the different methods. The three-line plots (from left to right) represent the performance of all spectra, spectra can be annotated by spectral matching, and spectra cannot be annotated by spectral matching. **b**. C1-C2 are the correct structures of the synthesized compounds, accompanied by the ranking given by DeepMASS. The predicted nearest neighbors are generally share the similar molecular skeleton and functional groups. This information is effective to locate the chemical space of the compounds to annotate. C1 and its nearest neighbor have clear common fragments, while the situations of C2 and C3 are less apparent. However, the Word2vec model is designed to learn non-intuitive correlations between fragments, which can be valuable in linking these correlations to the overall structures. **c**. DeepMASS ranks three isomeric candidates ahead of the correct structure of C2, which are similar to a certain extent. Despite not being the exact match, these candidates still provide valuable reference points for inferring the true structure.

Secondly, in order to evaluate the potential of DeepMASS to identify unknown compounds which have never discovered before. The restricted condition is the possible structures (candidates) are predicted and have been added to the database. A promising application identifying natural product derivatives. As a forward-looking example, three compounds beyond current libraries were synthesized. These derivatives of indole were synthesized strictly according to the protocol of the literature in which they were published[57]. Therefore, their structures cannot be identified by any spectral matching methods. Hence, we measured their mass spectra with ESI-FTMS instrument, and input the results to DeepMASS. The identification results are shown in **Fig. 3b**, accompanied with the nearest neighbors and number of candidates. Corrected structure of C1 and C3 are ranked at the first and second position, respectively (**Fig. 3b**). The correct structure C2 is ranked at the fourth position. Three isomeric candidates, which have similar substructures, are ranked before C2. Their corresponding structural similarities between these candidates and the correct structure calculated by MACCS key are 0.328, 0.719 and 0.483, respectively. Particularly, the second candidate share the very similar molecular skeleton. The results indicate DeepMASS can provide valuable reference to identify the unknown compounds (**Fig. 3c)**.

### GUI software and user instruction

We developed DeepMASS as a cross-platform GUI software tool, with the front end implemented by the QT framework and the back end implemented by the Python programming language. The dependent data of the public version was accompanied with the software, and we also provided the detailed manual for prepare the appropriate data format with the NIST and customized library. The software has been tested on Windows 11, Ubuntu and MacOS. As depicted in **Fig. 4**, the user interface is straightforward and user-friendly. For the sake of convenience and broad applicability, upon opening the software, the semantic model and reference dataset are automatically loaded. The reference dataset in the public version exclusively encompasses spectra from GNPS, but we offer scripts to assist users in incorporating their own in-house data and NIST data if they possess the necessary permissions.

**Fig. 4.**
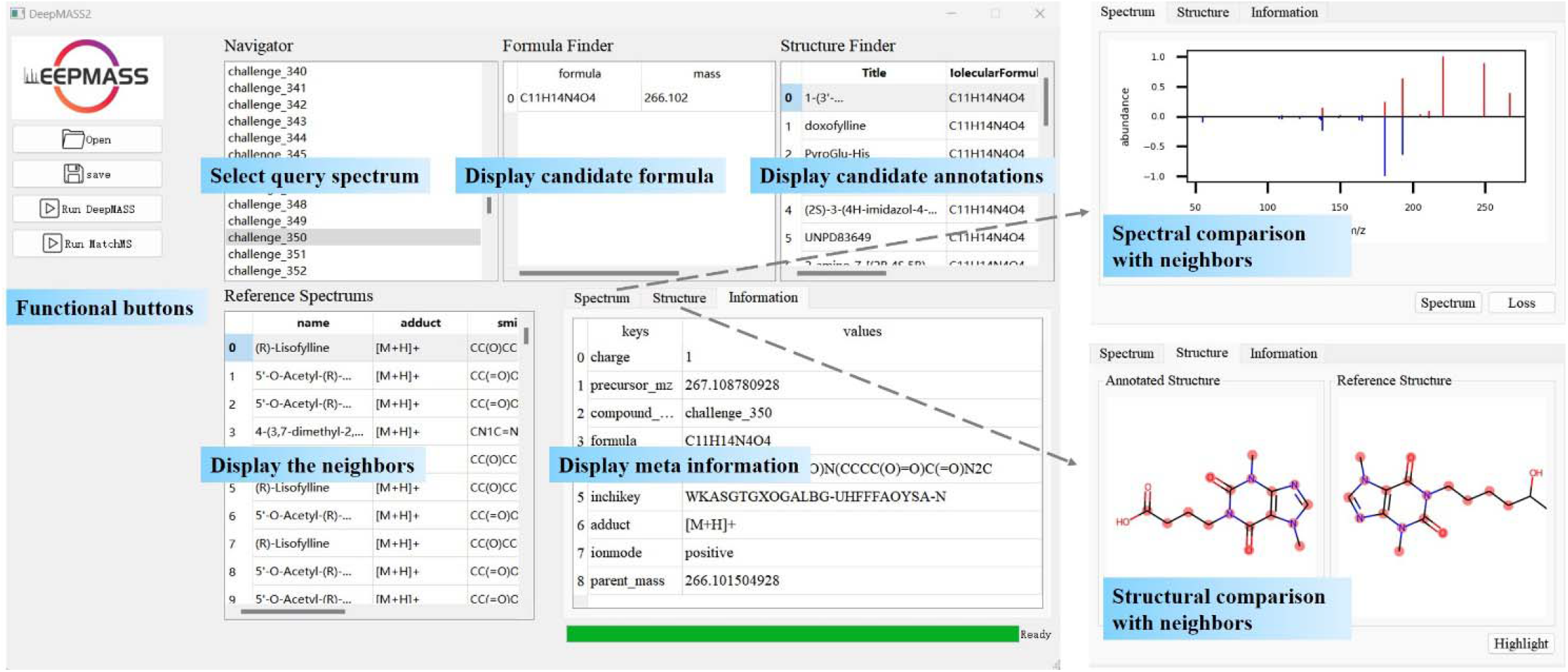
Software functionality overview. The software loads spectra via a mgf file and offers two modes: spectrum matching using MatchMS and in silico annotation with DeepMASS. It includes user-friendly widgets for displaying spectra, metadata, and predicted results. Additionally, it provides essential decision-making information to assess confidence in obtaining accurate results.

Unidentified spectra are uploaded in batches as *mgf* format files. DeepMASS autonomously recognizes and obtains the metadata for each spectrum. For instance, the ion mode dictates which model and reference dataset should be used for annotation. If the formula is included in the metadata, DeepMASS retrieves candidates with that specific formula; otherwise, it retrieves candidates based on the parent mass. The annotation results can be saved as an Excel file by clicking the *save* button.

The display panels consist of three sections: *spectrum, structure*, and *information*. Within the *spectrum* panel, a visual comparison is presented between the unidentified spectrum and the reference spectrum. Users have the option to choose the reference spectrum for comparison by clicking the corresponding item in the reference spectrum list widget. This feature helps users observe how DeepMASS reaches its annotation decisions and assesses the reliability of the annotations. Within the *structure* panel, a comparison is made between the structure of the reference compound and the candidate annotation. Through highlighting shared substructures, users can access additional explanatory information about the annotation. The *information* panel is used to show the meta information of the input spectrum. In general, DeepMASS is a user-friendly and accessible software tool for compound annotation that is designed for users who may not have programming skills.

### Case study for tomato ripening processes

To explore the practical utility of DeepMASS, we conducted a metabolomic analysis of the *Qianxi* cultivar cherry tomato during ripening processes (**Fig. 5a**). For the DeepMASS pipelines, data preprocessing and MS/MS extraction were conducted through MS-DIAL[35]. In total, 21,601 features assigned with MS/MS were obtained for the positive ion mode, and 18,846 features were obtained for the negative ion mode. DeepMASS successfully annotated 20,140 features from the positive mode data and 17,776 features from the negative mode data, accounting for 93.2% and 94.3% of the total, respectively. In comparison, TraceFinder, a commercial software used in the contrasting pipeline, annotated 21.7% of the features in the positive ion mode and 14.0% in the negative ion mode with library matching (**Fig. 5b**).

**Fig. 5.**
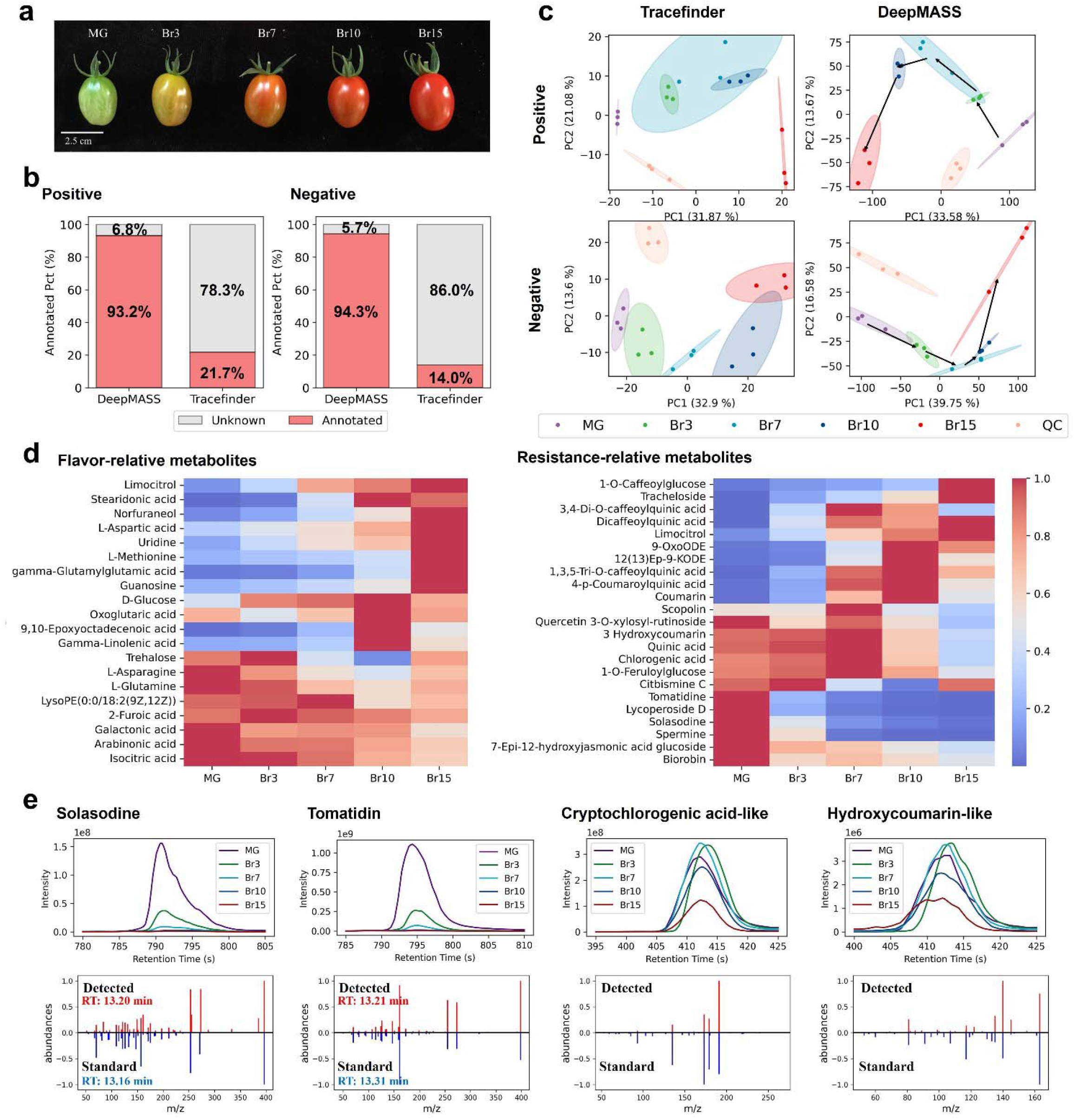
Tomato ripening processes analysis. a. Assorted ripening stages of tomato samples. b. Comparative analysis of annotated feature percentages by DeepMASS and TraceFinder. c. PCA scatter plot of samples with annotated compounds by DeepMASS and TraceFinder. d. Heatmap of the relative abundances of the identified metabolites annotated in association with various tomato ripening stages. e. XIC and MS/MS comparison of four identified metabolites.

We conducted a statistical analysis to compare the changes in the metabolomes from different ripening stages of *Qianxi* cherry tomatoes, using DeepMASS and TraceFinder annotations. Principal component analysis (PCA) was employed to assess metabolite differences among ripening stages. As shown in the score plot (**Fig. 5c**), based on the DeepMASS results, it was easy to distinguish fruit samples from different ripening stages completely, whereas it is less easy to effectively distinguish sample clusters from TraceFinder. Additionally, a continuous trend was observed in the principal component space with fruit ripening. This indicates that the additional metabolites annotated by DeepMASS were valuable in elucidating the metabolic changes during the ripening process of *Qianxi* cherry tomatoes.

Subsequently, we performed partial least squares discriminant analysis (PLS-DA) to identify biomarkers within the DeepMASS annotations, focusing on those with variable importance (VIP) scores exceeding 2 (**Table S4**). As the results, many of the identified metabolites have previously been linked to attributes such as resistance and fruit quality[58–60]. We then examined the variation trends of these metabolites across different ripening stages. Notably, we observed a gradual enrichment in the abundance of metabolites related to flavor and nutrients, including glucose, norfuraneol, glutamine, citric acid, and aspartic acid (**Fig. 5d**). These trends align with known variations in flavor and nutrient-related metabolites during ripening. We observed a decline in metabolites related to resistance but undesirable for consumers, such as solasodine, tomatidine, cryptochlorogenic acid-like compounds, and hydroxycoumarin-like compounds, especially in the later ripening stages. This pattern is consistent with the observed trends in the cherry tomato cultivar. To validate these findings, we compared the detected MS/MS with with standard substances. Additionally, we extracted the XIC of these metabolites and confirmed the trends with their chromatographic peaks. Two biomarkers are verified with retention times simultaneously, while for cryptochlorogenic acid-like and hydroxycoumarin-like compounds, DeepMASS cannot distinguish the positional isomers, preventing confirmation by retention times (**Fig. 5e**). Overall, the findings underscore the potential of the DeepMASS in obtaining a more comprehensive and accurate metabolome. This enhanced capability significantly improves the discovery potential for metabolites of interest.

## DISCUSSION

In this study, we introduce DeepMASS v2 as an enhanced iteration of our previous work[27]. The primary advancement is the strategy changing for finding the neighbor structures. With the incorporation of unsupervised semantic similarity model of the mass spectral language, the number of spectra for training increase from 752 to approximate 780,000, which is a three orders of magnitude improvement. The enhanced DeepMASS boasts the capability exhibits strong scalability, thus expanding its potential applications to various chemobiological fields.

Our positioning of DeepMASS is a competitive and collaborative program of SIRIUS and MS-Finder in MS/MS-based compound identification. They share common objective of structural annotation for compounds, irrespective of the availability of corresponding spectra. This strategic integration addresses the limitations in coverage observed in standard spectral databases, ensuring a more comprehensive approach to compound identification. This goal makes them distinguished from enhanced spectral matching software such as MatchMS or MS-Entropy, as well as from spectral embedding algorithms for numerical representation and spectral network establishment, like MS2DeepScore or Spec2Vec.

With the CASMI and in-house benchmark datasets, DeepMASS achieves higher annotation accuracy than the competitive software. Considering that some of the test spectra have corresponding reference spectra in the spectral database, we evaluate the performance separately for spectra that are within the database and those that are not. As expected, both DeepMASS and the competitive software exhibit significantly better performance when using spectra already present in the database. However, only DeepMASS demonstrates comparable performance when compared to the spectral matching strategy. This superiority is significant because, in practical applications, we often do not have prior knowledge of whether the spectra to be annotated are present in the database or not. When researchers use spectral matching and in silico annotation independently, the outcomes can diverge due to their distinct coverage of the chemical space. While spectral matching might excel in accuracy for common compounds, it can introduce bias and hinder the discovery of new compounds. DeepMASS circumvents this predicament by more effectively leveraging the reference database compared with the other in silico annotation software. When the compound is already present in the database, it serves as the optimal choice for localizing the chemical space, and the other neighbors can complement and enhance its performance. This approach yields results that surpass those obtained by using reference spectra in isolation.

On the other hand, in cases where the spectra lack corresponding compounds within the reference database, the predicted neighbors based on spectral similarity with the unknown spectra play a pivotal role in capturing the chemical space and evaluating potential candidates. Since spectral similarity is a reflection of structural similarity, this prediction is logically justified. Moreover, given that diverse functional metabolites typically stem from a restricted set of initial metabolites through biochemical reactions, there exists a reasonable probability that reference neighbors might lie upstream or downstream of the unknown metabolite within metabolic pathways. This circumstance enhances the likelihood of accurate annotations.

To evaluate the application value in an actual metabolomics study, we conducted a metabolomics analysis of the *Qianxi* cultivar cherry tomato during ripening processes and found that DeepMASS significantly increased the utilization rate of raw data and effectively reduced data bias in subsequent analysis. In addition, from the annotated results, we also found a series of metabolites that might contribute to tomato fruit flavor and commercial quality trait formation during ripening. In summary, metabolomic analysis of these tomato fruit samples confirmed that DeepMASS not only substantially enhanced the extraction of features and the annotation of metabolites from raw mass spectral data but also did so with high reliability. This underscores its utility as a potent tool for the discovery of new metabolite biomarkers in various biological contexts.

Looking ahead, DeepMASS shows promise in effectively identifying undiscovered compounds, on condition that the possible structures are predicted and added into the database. A promising showcase of identifying natural product derivatives is presented in the supplementary information. The limitation of the current version of DeepMASS is its strong dependence on both the quantity and quality of spectra within the reference database. It is evident from the varying performance observed when NIST spectra are included as reference data versus when they are not. This reliance is crucial for identifying reasonable and accurate neighbor compounds that assist in pinpointing the chemical space of the unknown metabolite. The expansion of spectral data further enhances the precision of the connection between structural similarity and semantic similarity within the mass spectral language. Overall, DeepMASS demonstrates commendable annotation performance in both benchmark evaluations and real-world data analysis. It offers meaningful and valuable metabolite annotations, features user-friendly and extensible software, and holds promise for even greater potential with the incorporation of more reliable data in the future.

## METHODS

### Semantic model training

Three distinct datasets were utilized to train the word2vec-based semantic model. Initially, the open-source GNPS dataset, acquired as of June 1st, 2022, comprised 495,810 MS/MS spectra. Second, the commercial NIST 20 software contributed 1,026,506 MS/MS spectra. Last, our in-house dataset added an additional 23,610 MS/MS spectra. A publicly available version excluding the NIST 20 training set is provided due to commercial licensing constraints.

The spectra underwent a cleansing and correction process facilitated by the MatchMS package[61], including the following procedures: (1) extract structural information (SMILES, InChI, InChIKey) from the metadata, and discard entries for which this information could not be derived; (2) extract information related to precursor type (adduct, charge) from the metadata, retaining only the spectra with adduct type of M+H, M+Na, M+K, and M+NH_4_ for positive mode and M-H and M+Cl for negative mode; (3) exclude spectra with fewer than five fragment ions; (4) normalize peak intensities to a range of 0 to 1; and (5) to avoid any overlap in data, eliminate all spectra with cosine similarity greater than 0.95 compared to any spectrum in the CASMI datasets. Consequently, 600,289 positive spectra derived from 43,653 unique compounds and 182,323 distinct negative spectra from 20,883 unique compounds were retained.

After processing, spectra were transformed into documents. For this, each peak was represented by a word containing its position up to a specified decimal precision (). In all presented outcomes, a binning approach involving two decimal places was applied, meaning that a peak at m/z 200.445 was represented as the word “peak@200.45“. Alongside all the peaks within a spectrum, neutral losses ranging from 5.0 to 200.0 Da were also included as. The calculation of neutral losses involved the precursor m/z subtracted from the peak m/z. The compilation of all the generated peak and loss words constitutes what is referred to here as a document. Two semantic models were separately trained using the Spec2Vec package[28] for spectra in both positive and negative modes. The hyperparameters were configured as follows: (1) word vector dimension = 300, (2) negative sampling (negative = 5), (3) initial learning rate = 0.025, (4) learning rate decay per epoch = 0.00025, and (5) number of epochs = 30. During the parameter optimization process, 10% of the dataset was separated from the training set to form the validation set. However, when training the final model, the entire training set was utilized. After training, the semantic information is expected to be captured through the Word2Vec model.

### Neighbor compound searching

Using our trained semantic model, we embedded all spectra from the training set into vectors. This process resulted in two matrices: a 600,289 × 300 matrix for positive mode and a 182,323 × 300 matrix for negative mode. Each row in these matrices represents an embedded vector, and the similarity between vectors correlates with the structural similarity of the corresponding molecules. When presented with an unknown spectrum, it is embedded into a vector with the same model. Previous studies on Spec2Vec have demonstrated that the distance between the semantic model embedded vectors effectively reflects the structural similarity of corresponding metabolites[28]. Here, our objective is to efficiently compare the embedded vector of the query spectrum with pre-built matrices.

Therefore, we employed Hierarchical Navigable Small-World (HNSW) graphs. HNSW is an Approximate Nearest Neighbor Search (ANNS) method based on hierarchical graph structures and graph traversal techniques. HNSW indices do not require training with data. The constructional parameters are *M* = 64 and *ef_construction* = 800, where *M* defines the maximum number of outgoing connections in the graph, and *ef_construction* defines a construction time/accuracy trade-off. Essentially, the HNSW index serves as an efficient data structure, organizing hierarchically multi-layer graphs to store, organize, and calculate distances between vectors. This approach significantly improves the speed of ANNS. ANNS method is applied for rapid retrieval of the Top 300 vectors with the least distance. The structures associated with the retrieved vectors are considered “neighbor compounds”, indicating that they likely share similar structures with the unknown.

### Candidate retrieval and ranking

A typical metabolomic data processing workflow utilizing comprehensive software like XCMS[31], MZMine[32], OpenMS[33], KPIC2[34] or MS-DIAL[35] provides quantitative features along with corresponding information such as precursor m/z, ion type, charge, and isotopic pattern for unknowns. This information enables the users annotate the molecular formulas using external software tools like BUDDY[25] or MIST-CF[26]. Therefore, DeepMASS encourages users to input unknown spectra along with assigned formulas, allowing it to retrieve candidates from the structural bio database based on the provided formula. In cases where uploaded spectra lack this information, DeepMASS requires precursor m/z, ion type, charge, and isotopic pattern. It then infers the monoisotopic mass based on these parameters and retrieves and evaluates potential candidates with a monoisotopic mass difference within a set threshold (defaulting to 10 ppm).

DeepMASS evaluates and ranks candidates by considering the relative location between the encompassed region of “neighbor compounds” and the candidates within the chemical space. The position of compounds in the chemical space is estimated using Morgan fingerprints, and the distance between paired compounds is determined through the *Dice* distance calculated from their fingerprint vectors.

To elaborate, for each candidate, DeepMASS scans the “neighbor compounds”, identifying the nearest 20 compounds. The algorithm then calculates the cumulative distance between these selected “neighbor compounds” and the candidate, using this summed distance as the ranking score. The mathematical expression is:

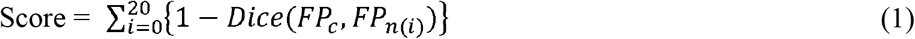

where *FP*_*c*_ represents the fingerprints of the candidate compound, *FP*_*n*(*i*)_ represents the fingerprints of the *i*_*th*_ “neighbor compounds”, and Dice refers to the Dice distance. From the equation, it can be seen that the candidates with higher scores are closer to the predicted structurally similar neighbors.

According to the equation, candidates which are closer to the predict neighbor compounds will receive higher scores. In fact, it’s not necessary for all predicted compounds to be highly related to the unknown substance. A subset of neighbor compounds showing structural similarities with the unknown substance is adequate to map the chemical space, thereby allowing the correct structure to receive a significantly higher score than others. This feature also means that the prediction of neighbors does not have to be exceptionally accurate. A few false positive predictions do not significantly impact the identification results.

### CASMI benchmarking data

We utilized CASMI 2014, 2016, and 2022 as benchmarking data for our evaluation. MS/MS peak lists for CASMI 2014 and 2016 were readily available on the official website. In the case of CASMI 2022, raw files of *mzML* format are provided, we directly extracted MS/MS data from the raw files using provided retention time and precursor m/z information. CASMI 2017 was excluded due to its utilization of MS^E^ mode, as all the spectra of the reference dataset are from DDA mode. We assumed that correct molecular formulas were annotated and could be directly employed for candidate retrieval in the software utilized for comparison.

### Natural product benchmarking data

Natural products were purchased from MedChemExpress company (MCE). Each reagent is taken at 2 μL and diluted with MeOH/MeCN/H2O (V:V:V=2:2:1) to a stock solution concentration of 50 μg/mL, with the remaining reagents stored at -80°C. For every group of 50 reagents, take 5 μL from each stock solution and mix them together. Create a table to indicate the types of compounds contained in each sample and ensure that no compounds with the same molecular weight are included within the same sample.

Mixtures were separated by an ACQUITY UPLC™ BEH C18 column (Waters Ltd., MA, USA) (100 × 2.1 mm, 2.6 µm), and were analyzed by a Vanquish UHPLC system coupled with a Q-Exactive HF-X orbitrap mass spectrometer (Thermo Fisher Scientific Ltd., CA, USA), equipped with a HESI probe. The injection volumn was set at 1 µL, and the flow rate was set at 300 μL/min. Solvents for the mobile phase were 0.05% formic acid in acetonitrile (A) and 0.1% formic acid in water (B). The gradient elution was: 0-25 min, linear gradient 30 %-100 % A, and then the column was washed with 100 % B and 100 % A. Data in the mass range of m/z 80-1200 were acquired with data-dependent MS/MS acquisition. The full scan and fragment spectra were collected at resolutions of 70,000 and 17,500, respectively. The exported raw files of the platform are converted into *mzML* with MSConvert. We extracted MS/MS data from the mzML files with PyOpenMS[62]. If more than two MS/MS were obtained, the one with the highest precursor intensity was selected.

### Competitive software

To assess the performance of DeepMASS, a comparative analysis was conducted using two renowned state-of-the-art in silico metabolite annotation software tools: MS-Finder (version 3.52) and SIRIUS (version 5.8.1). The parameter settings were as follows: SIRIUS, *MS2 mass accuracy* of 10 ppm, *Database* of ‘*Custom’, Use heuristic above m/z* of *300*; MF-Finder, *MS2 mass tolerance* of 0.01 Da, *Tree depth* of 2, *Database* of ‘*User defined database*’. Both DeepMASS and the comparative software employed spectra in *msp* format, the chemical formulas were considered known. SIRIUS applied the CSI:FingerID algorithm, while MS-Finder utilized in silico fragmentation algorithm to prioritize candidate rankings. The compound databases utilized by SIRIUS and MS-Finder were the same as those used by DeepMASS, through custom uploads via their parameter setting functions. A noteworthy aspect is that both SIRIUS and MS-Finder have already integrated all GNPS data into their training sets. Consequently, a portion of the CASMI data could be considered exposed to their models.

### Tomato ripening study protocol

Cherry tomato (cultivar *Qianxi*) was used for the transgenic experiments. The plant materials were planted in the greenhouse at the AGIS experimental station in Shenzhen, China (114°30’3.63” E, 22°36’4.66” N). All seedlings were grown in a commercial nursery for 30-40 days at 28°C and then transplanted to fields. Three to five fruits per plant were picked at selected ripe stages and immediately frozen in liquid nitrogen. Then, the frozen tissues were ground to fine powders with an IKA A11 machine (IKA Works, Inc., NC, USA) and stored in a -80 □ freezer for metabolomic analysis.

For nontarget metabolomic analysis, 100 mg of tomato powder for each sample was extracted with 1 mL of extraction buffer (MeOH:H2O = 80:20, v/v, containing 0.1% formic acid, with 100 ng/mL of d_5_-coumarin, d_5_-L-phenylalanine, and d_3_-homovanillic acid as internal standards (Cambridge Isotope Laboratories, Inc., MA, USA)), followed by a 30 min ultrasonic treatment. The supernatants were pipetted and dried in a LABCONCO CentriVap vacuum centrifugal concentrator (We Brothers Scientific (Pvt.) Ltd., MO, USA) and resuspended in 100 µL of reconstitution buffer (MeOH:H2O = 80:20, v/v). A 10 µL aliquot of the supernatant from each sample was pipetted and mixed as QC samples.

Then, extracts were analyzed by a U3000 UHPLC system coupled with a Q Exactive orbitrap mass spectrometer (Thermo Fisher Scientific Ltd., CA, USA) equipped with a HESI probe under both positive and negative modes. Extracts were separated by an ACQUITY UPLC™ BEH C18 column (Waters Ltd., MA, USA) (150 × 2.1 mm, 2.6 µm). The injection volume was set at 1 µL, and the flow rate was set at 300 μL/min. Solvents for the mobile phase were 0.05% formic acid in acetonitrile (A) and 0.1% formic acid in water (B). The elution gradient was as follows: 0-25 min, linear gradient 40%-100% A, followed by column washing with 100% B and 100% A. Data with mass ranges of m/z 80-1200 and m/z 80-1200 were acquired in both positive ion mode and negative ion mode separately, with data-dependent MS/MS acquisition. The full scan and fragment spectra were collected with resolutions of 70,000 and 17,500, respectively.

For spectral matching pipelines, TraceFinder analysis software (version 4.1) was used to analyze the acquired MS/MS data and process feature annotation with commercial and public databases. The data processing parameter settings were as follows: *peak width* from 10 to 45 s, *mzwid* of 0.015, *minfrac* of 0.5, *bw* of 5, and signal/noise threshold of 6. Then, compounds were annotated based on our in-house database via high-resolution MS-associated methods. Accurate high-resolution m/z values and MS/MS fragmentation patterns were used to identify metabolites, with a mass accuracy of 2 ppm and RT window of 20 s. Then, the annotated metabolites were checked and confirmed by comparing retention times, fragment patterns and isotope peaks manually. The library matching score threshold was set as 30.

For the DeepMASS pipelines, data preprocessing and MS/MS extraction were conducted through MS-DIAL[35]. The data processing parameter settings were as follows: *peak height* of 20000, *mass slice* of 0.1 Da, *alignment retention time tolerance* of 0.05 min. Then, MS/MS files were exported and fed to DeepMASS software. Subsequently, the MS/MS files were exported and inputted into the DeepMASS software. The sole parameter setting adjusted was the candidate retrieving mass tolerance, which was set to 10 ppm. The retrieval range defaulted to the entire database, while the structural annotation processing is non-parametric.

## DATA AVAILABILITY

Spectral data from GNPS are available at the official website (https://gnps.ucsd.edu/ProteoSAFe/static/gnps-splash.jsp); Spectral data from NIST 20 are commercially available and can be purchased from multiple vendors. CASMI benchmarking data are available at the CASMI website (http://casmi-contest.org); Raw *mzML* files of Natural product benchmarking data, the preprocessed spectral data of CASMI and natural product, accompanied with the annotated results of DeepMASS, SIRIUS and MS-Finder are available at https://github.com/hcji/DeepMASS2_Data_Processing.

## CODE AVAILABILITY

The software and source code for data cleaning, model training and performance evaluation of DeepMASS have been released under the AGPL 3.0 license at https://github.com/hcji/DeepMASS2_GUI. The data processing and performance evaluation scripts are available at https://github.com/hcji/DeepMASS2_Data_Processing.

## ACKNOWLEDGMENTS

This work is financially supported by National Key R&D Program of China (grant. 2023YFA0915800) and the National Natural Science Foundation of China (Grant No. 22103036) and. It was also supported by Innovation Program of Chinese Academy of Agricultural Sciences and the Elite Young Scientists Program of CAAS.

## AUTHOR CONTRIBUTIONS

Conceptualization, H.J.; methodology, H.J.; formal analysis: H.J.; investigation: H.J., R.D.; resources: R.D.; Y.L.; J.Y.; experiment: S.F.; data curation: H.J., Q.D.; statistics: R.D.; writing original draft preparation: H.J., R.D.; review and editing, H.J., J.Y.; visualization: H.J., R.D.; supervision: J.Y.; project administration, J.Y.

## REFERNCES

1. Chen C-J, Lee D-Y, Yu J, et al. Recent advances in LC-MS-based metabolomics for clinical biomarker discovery. Mass Spectrom. Rev. 2023; 42:2349–2378

2. Shen S, Zhan C, Yang C, et al. Metabolomics-centered mining of plant metabolic diversity and function: Past decade and future perspectives. Mol. Plant 2023; 16:43–63

3. Herrero M, Simó C, García-Cañas V, et al. Foodomics: MS-based strategies in modern food science and nutrition. Mass Spectrom. Rev. 2012; 31:49–69

4. Maitre L, Bustamante M, Hernández-Ferrer C, et al. Multi-omics signatures of the human early life exposome. Nat. Commun. 2022; 13:7024

5. Bazsó FL, Ozohanics O, Schlosser G, et al. Quantitative Comparison of Tandem Mass Spectra Obtained on Various Instruments. J. Am. Soc. Mass Spectrom. 2016; 27:1357–1365

6. Li Y, Kind T, Folz J, et al. Spectral entropy outperforms MS/MS dot product similarity for small-molecule compound identification. Nat. Methods 2021; 18:1524–1531

7. De Jonge NF, Louwen JJR, Chekmeneva E, et al. MS2Query: reliable and scalable MS2 mass spectra-based analogue search. Nat. Commun. 2023; 14:1752

8. Yang Q, Ji H, Xu Z, et al. Ultra-fast and accurate electron ionization mass spectrum matching for compound identification with million-scale in-silico library. Nat. Commun. 2023; 14:3722

9. Da Silva RR, Dorrestein PC, Quinn RA. Illuminating the dark matter in metabolomics. Proc. Natl. Acad. Sci. 2015; 112:12549–12550

10. Wang M, Carver JJ, Phelan VV, et al. Sharing and community curation of mass spectrometry data with Global Natural Products Social Molecular Networking. Nat. Biotechnol. 2016; 34:828–837

11. van der Hooft JJJ, Wandy J, Barrett MP, et al. Topic modeling for untargeted substructure exploration in metabolomics. Proc. Natl. Acad. Sci. 2016; 113:13738–13743

12. Morehouse NJ, Clark TN, McMann EJ, et al. Annotation of natural product compound families using molecular networking topology and structural similarity fingerprinting. Nat. Commun. 2023; 14:308

13. Cai Y, Zhou Z, Zhu Z-J. Advanced analytical and informatic strategies for metabolite annotation in untargeted metabolomics. TrAC Trends Anal. Chem. 2023; 158:116903

14. Hu G, Qiu M. Machine learning-assisted structure annotation of natural products based on MS and NMR data. Nat. Prod. Rep. 2023; 40:1735–1753

15. Dührkop K, Shen H, Meusel M, et al. Searching molecular structure databases with tandem mass spectra using CSI:FingerID. Proc. Natl. Acad. Sci. 2015; 112:12580–12585

16. Dührkop K, Fleischauer M, Ludwig M, et al. SIRIUS 4: a rapid tool for turning tandem mass spectra into metabolite structure information. Nat. Methods 2019; 16:299–302

17. Tsugawa H, Kind T, Nakabayashi R, et al. Hydrogen Rearrangement Rules: Computational MS/MS Fragmentation and Structure Elucidation Using MS-FINDER Software. Anal. Chem. 2016; 88:7946–7958

18. Wang F, Liigand J, Tian S, et al. CFM-ID 4.0: More Accurate ESI-MS/MS Spectral Prediction and Compound Identification. Anal. Chem. 2021; 93:11692–11700

19. Lai Z, Tsugawa H, Wohlgemuth G, et al. Identifying metabolites by integrating metabolome databases with mass spectrometry cheminformatics. Nat. Methods 2018; 15:53–56

20. Ji H, Deng H, Lu H, et al. Predicting a Molecular Fingerprint from an Electron Ionization Mass Spectrum with Deep Neural Networks. Anal. Chem. 2020; 92:8649–8653

21. Wang L, Ye H, Sun D, et al. Metabolic Pathway Extension Approach for Metabolomic Biomarker Identification. Anal. Chem. 2017; 89:1229–1237

22. Shen X, Wang R, Xiong X, et al. Metabolic reaction network-based recursive metabolite annotation for untargeted metabolomics. Nat. Commun. 2019; 10:1516

23. Zhou Z, Luo M, Zhang H, et al. Metabolite annotation from knowns to unknowns through knowledge-guided multi-layer metabolic networking. Nat. Commun. 2022; 13:6656

24. Kim S, Thiessen PA, Bolton EE, et al. PUG-SOAP and PUG-REST: Web services for programmatic access to chemical information in PubChem. Nucleic Acids Res. 2015; 43:W605–W611

25. Xing S, Shen S, Xu B, et al. BUDDY: molecular formula discovery via bottom-up MS/MS interrogation. Nat. Methods 2023; 20:881–890

26. Goldman S, Xin J, Provenzano J, et al. MIST-CF: Chemical Formula Inference from Tandem Mass Spectra. J. Chem. Inf. Model. 2023;

27. Ji H, Xu Y, Lu H, et al. Deep MS/MS-Aided Structural-Similarity Scoring for Unknown Metabolite Identification. Anal. Chem. 2019; 91:5629–5637

28. Huber F, Ridder L, Verhoeven S, et al. Spec2Vec: Improved mass spectral similarity scoring through learning of structural relationships. PLOS Comput. Biol. 2021; 17:e1008724

29. Huber F, van der Burg S, van der Hooft JJJ, et al. MS2DeepScore: a novel deep learning similarity measure to compare tandem mass spectra. J. Cheminformatics 2021; 13:84

30. Guo H, Xue K, Sun H, et al. Contrastive Learning-Based Embedder for the Representation of Tandem Mass Spectra. Anal. Chem. 2023; 95:7888–7896

31. Forsberg EM, Huan T, Rinehart D, et al. Data processing, multi-omic pathway mapping, and metabolite activity analysis using XCMS Online. Nat. Protoc. 2018; 13:633–651

32. Schmid R, Heuckeroth S, Korf A, et al. Integrative analysis of multimodal mass spectrometry data in MZmine 3. Nat. Biotechnol. 2023; 41:447–449

33. Pfeuffer J, Bielow C, Wein S, et al. OpenMS 3 enables reproducible analysis of large-scale mass spectrometry data. Nat. Methods 2024; 1–3

34. Ji H, Zeng F, Xu Y, et al. KPIC2: An Effective Framework for Mass Spectrometry-Based Metabolomics Using Pure Ion Chromatograms. Anal. Chem. 2017; 89:7631–7640

35. Tsugawa H, Cajka T, Kind T, et al. MS-DIAL: Data-independent MS/MS deconvolution for comprehensive metabolome analysis. Nat. Methods 2015; 12:523–526

36. Malkov YA, Yashunin DA. Efficient and Robust Approximate Nearest Neighbor Search Using Hierarchical Navigable Small World Graphs. IEEE Trans. Pattern Anal. Mach. Intell. 2020; 42:824–836

37. Hastings J, Owen G, Dekker A, et al. ChEBI in 2016: Improved services and an expanding collection of metabolites. Nucleic Acids Res. 2016; 44:D1214–D1219

38. Barupal DK, Fiehn O. Generating the Blood Exposome Database Using a Comprehensive Text Mining and Database Fusion Approach. Environ. Health Perspect. 2019; 127:097008

39. Knox C, Wilson M, Klinger CM, et al. DrugBank 6.0: the DrugBank Knowledgebase for 2024. Nucleic Acids Res. 2023; gkad976

40. Sajed T, Marcu A, Ramirez M, et al. ECMDB 2.0: A richer resource for understanding the biochemistry of E. coli. Nucleic Acids Res. 2016; 44:D495–D501

41. Naveja JJ, Rico-Hidalgo MP, Medina-Franco JL. Analysis of a large food chemical database: chemical space, diversity, and complexity. 2018;

42. Wishart DS, Guo A, Oler E, et al. HMDB 5.0: the Human Metabolome Database for 2022. Nucleic Acids Res. 2022; 50:D622–D631

43. Kanehisa M, Furumichi M, Tanabe M, et al. KEGG: New perspectives on genomes, pathways, diseases and drugs. Nucleic Acids Res. 2017; 45:D353–D361

44. Ntie-Kang F, Telukunta KK, Döring K, et al. NANPDB: A Resource for Natural Products from Northern African Sources. J. Nat. Prod. 2017; 80:2067–2076

45. van Santen JA, Jacob G, Singh AL, et al. The Natural Products Atlas: An Open Access Knowledge Base for Microbial Natural Products Discovery. ACS Cent. Sci. 2019; 5:1824–1833

46. Tian Z, Hu X, Xu Y, et al. PMhub 1.0: a comprehensive plant metabolome database. Nucleic Acids Res. 2023; gkad811

47. Hawkins C, Ginzburg D, Zhao K, et al. Plant Metabolic Network 15: A resource of genome-wide metabolism databases for 126 plants and algae. J. Integr. Plant Biol. 2021; 63:1888–1905

48. Frolkis A, Knox C, Lim E, et al. SMPDB: The Small Molecule Pathway Database. Nucleic Acids Res. 2010; 38:D480–D487

49. Williams AJ, Grulke CM, Edwards J, et al. The CompTox Chemistry Dashboard: a community data resource for environmental chemistry. J. Cheminformatics 2017; 9:61

50. Wishart D, Arndt D, Pon A, et al. T3DB: the toxic exposome database. Nucleic Acids Res. 2015; 43:D928–D934

51. Ru J, Li P, Wang J, et al. TCMSP: a database of systems pharmacology for drug discovery from herbal medicines. J. Cheminformatics 2014; 6:13

52. Ramirez-Gaona M, Marcu A, Pon A, et al. YMDB 2.0: a significantly expanded version of the yeast metabolome database. Nucleic Acids Res. 2017; 45:D440–D445

53. Nothias L-F, Petras D, Schmid R, et al. Feature-based molecular networking in the GNPS analysis environment. Nat. Methods 2020; 17:905–908

54. Simón-Manso Y, Lowenthal MS, Kilpatrick LE, et al. Metabolite Profiling of a NIST Standard Reference Material for Human Plasma (SRM 1950): GC-MS, LC-MS, NMR, and Clinical Laboratory Analyses, Libraries, and Web-Based Resources. Anal. Chem. 2013; 85:11725–11731

55. Kim HW, Wang M, Leber CA, et al. NPClassifier: A Deep Neural Network-Based Structural Classification Tool for Natural Products. J. Nat. Prod. 2021; 84:2795–2807

56. Becht E, McInnes L, Healy J, et al. Dimensionality reduction for visualizing single-cell data using UMAP. Nat. Biotechnol. 2019; 37:38–44

57. Gao J, He X-C, Liu Y-L, et al. Photoredox/Nickel Dual Catalysis-Enabled Cross-Dehydrogenative C–H Amination of Indoles with Unactivated Amine. Org. Lett. 2023; 25:7716–7720

58. Zhu G, Wang S, Huang Z, et al. Rewiring of the Fruit Metabolome in Tomato Breeding. Cell 2018; 172:249-261.e12

59. Tieman D, Zhu G, Resende MFR, et al. A chemical genetic roadmap to improved tomato flavor. Science 2017; 355:391–394

60. Klee HJ, Tieman DM. The genetics of fruit flavour preferences. Nat. Rev. Genet. 2018; 19:347–356

61. Huber F, Verhoeven S, Meijer C, et al. matchms - processing and similarity evaluation of mass spectrometry data. J. Open Source Softw. 2020; 5:2411

62. Röst HL, Schmitt U, Aebersold R, et al. pyOpenMS: A PythonLJbased interface to the OpenMS mass spectrometry algorithm library. PROTEOMICS 2014; 14:74–77

