## Supplementary material for "DeepMASS: Unknown Compound Annotation using Semantic Similarity of Mass Spectral Language and Chemical Space Localization": Fig. S1; Fig. S2 (from another version of this preprint: version 1, 2024-06-03)

### Supplementary Figures

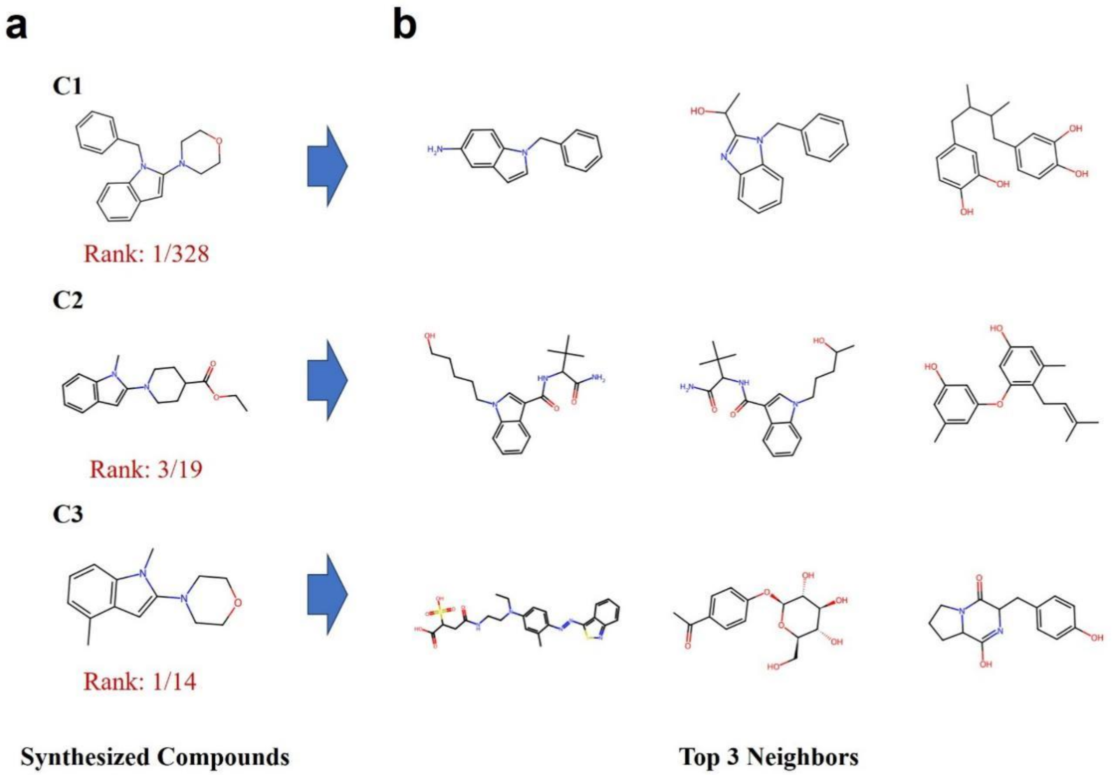


**Fig.S1** Structural identification of synthesized derivatives of indole. **a.** C1-C2 are the true two-dimensional structures of the derivatives. The ranking of the correct structure of each compound obtained by DeepMASS is labeled bellow the structures. **b**. The corresponding neighbor compounds predicted by the semantic model, which are used for encompass the chemical space of the unknown and scoring the candidates. These neighbor compounds are related to the correct structure, aiding in the identification process by DeepMASS.

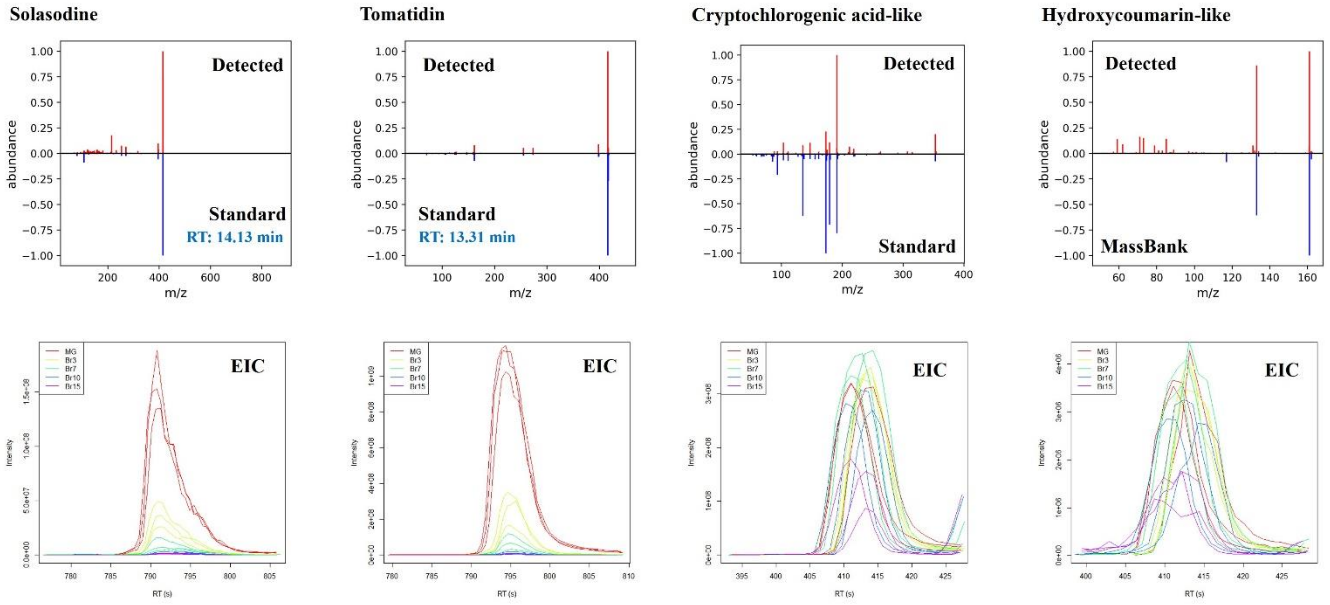


**Fig.S2** MS/MS and EICs of the metabolites decreased in abundance which are annotated as related to resistance. Solasodine, tomatidine and chlorogenic acid-like compounds are verified with standards, while hydroxycoumarin is verified with spectrum in MassBank.
